# Neural representations of speech production in neocortical and cerebellar regions

**DOI:** 10.64898/2026.01.30.702863

**Authors:** Sivan Jossinger, Bassel Arafat, Jorn Diedrichsen

## Abstract

Speech production requires the coordinated control of the supralaryngeal articulators (e.g., the lips, tongue, and soft palate) together with laryngeal control of phonation. While cortical sensorimotor regions are known to encode phonetic features, how the cerebellum represents speech and how cerebellar representations integrate within cortico-cerebellar circuits remain poorly understood. Using 7T functional MRI and representational similarity analysis, we examined neural representations of speech production during overt syllable production varying in place of articulation and voice onset time. We found reliable, syllable-specific activity patterns across both cortical and cerebellar speech regions. Ventral primary sensorimotor cortex distinguished syllables by place of articulation, whereas dorsal sensorimotor cortex showed sensitivity to voice onset time, consistent with a functional dissociation between articulatory and phonatory control. We also found secondary speech areas that encoded phonetic features. Specifically, the parietal operculum showed sensitivity to both place of articulation and voice onset time, positioning it as an integrative node within the speech motor network. In the cerebellum we found superior and inferior speech areas, which did not differ in their representational geometry. Surprisingly, the cerebellar syllable representations were most similar to those found in the parietal operculum rather than in primary motor cortex, with both regions encoding both phonetic features. These findings indicate that cerebellar speech representations are not solely determined by a single primary sensorimotor area but instead align more closely with the representations in the parietal operculum, suggesting that cerebellar speech areas contribute to the coordination of phonation and articulation.

## 1. Introduction

Speech is among the most complex motor activities humans perform, requiring the coordination of approximately 100 muscles across laryngeal, respiratory and oral motor systems. This coordination integrates two fundamental speech features: articulation, which shapes the configuration of the vocal tract, and phonation, the generation of voiced sound. Such intricate orchestration depends on a distributed neural network spanning both cortical and cerebellar regions (Guenther & Hickok, 2016). Prior imaging work has primarily characterized neocortical representations of speech features, yet how the cerebellum encodes these fundamental building blocks of speech remains poorly understood.

Cortical representations of speech motor control have been most thoroughly characterized in the primary sensorimotor cortex. The ventral sensorimotor cortex (vSM) projects directly to brainstem nuclei that innervate the vocal tract via multiple cranial nerves (Jürgens, 2002; Penfield & Boldrey, 1937). Neural populations in this region are selectively tuned to specific speech effectors, including the lips, tongue, jaw, and larynx, forming a somatotopic organization that mirrors the anatomy of the vocal tract (Bouchard et al., 2013; Carey et al., 2017). In contrast, a more dorsal region of the sensorimotor cortex (dSM) has been traditionally associated with trunk movements involved in voluntary breathing (Foerster, 1931; Penfield & Boldrey, 1937; but see Gordon et al., 2023). A recent study contrasting voiced and whispered speech reported greater activation in dSM during voiced speech, suggesting that this region in involved not only in respiration, but also in the control of phonation (Correia et al., 2020).

In addition to the primary sensorimotor cortex, the cerebellum plays a critical role in speech production (Ackermann & Brendel, 2016). Patients with cerebellar lesions often present with ataxic dysarthria, a speech motor control syndrome characterized by inaccurate articulation, harsh voice quality and disrupted speech timing, implicating the cerebellum in both aspects of speech motor control (Ackermann et al., 1997; Ackermann & Hertrich, 1997). Accordingly, kinematic and acoustic studies in cerebellar patients reveal deficits across both domains, including abnormal articulatory movement dynamics (Ackermann et al., 1995), as well as increased instability in pitch and voice quality during phonation (Ackermann & Ziegler, 1994; Ackermann, 1991).

Imaging studies show that overt speech activates two separate regions in each cerebellar hemisphere: a superior region in lobules VI and an inferior in lobules VIII (Ackermann, 2008; Bohland & Guenther, 2006; Correia et al., 2020). Prior works indicate that activity in these two regions may differ in the specific task demands that engage them (Bohland & Guenther, 2006; Chen & Desmond, 2005; Riecker et al., 2000). Moreover, speech deficits following cerebellar stroke most commonly occur after lesions in the territory of the superior cerebellar artery, which supplies the superior cerebellum (Ackermann et al., 1992). This raises the possibility that superior and inferior cerebellar speech regions support different aspects of speech processing.

Both cerebellar speech regions overlap with cerebellar motor territories implicated in tongue movement (Nettekoven et al., 2024; Saadon-Grosman et al., 2022). In the limb motor system, the cerebellar hand area is reciprocally connected with the hand representation in the sensorimotor cortex, forming a closed cortico-cerebellar loop (Kelly & Strick, 2003; Saadon-Grosman et al., 2022). By analogy, the cerebellar speech areas may be preferentially coupled with vSM, in which case they should exhibit articulatory representations similar to those observed in vSM. Alternatively, they may be preferentially connected with the dorsal sensorimotor area (dSM), predicting representations that primarily reflect phonatory control. A third possibility is that cerebellar speech areas resemble higher-order neocortical speech regions, such as the supplementary motor area (SMA) or speech-related regions in the temporoparietal junction.

To address this question, we used high-field (7T) fMRI while participants produced syllables that differed in place of articulation and voicing. We combined high-resolution imaging with representational similarity analysis (RSA) to characterize how specific speech features are encoded in cortical and cerebellar activity patterns. Consistent with previous work (Bouchard et al., 2013), activity patterns in vSM primarily distinguished between places of articulation. In contrast, representational geometries in dSM differentiated between voiced and voiceless consonants, in line with the proposed role of this region in phonation (Correia et al., 2020). Secondary sensorimotor speech areas outside primary motor cortex showed a mixture of these two representations. We then asked whether cerebellar speech representations resemble those observed in cortical speech regions.

## 2. Methods

### Participants

Twelve neurotypical adults were recruited for this study (6 females, 18-29 years [mean ± SD = 23.3 ± 3.6]). All participants were right handed as estimated by the Edinburgh handedness inventory (90.7 ± 15.1; Oldfield, 1971), native-level English speakers, and had no history of speech impairment or a neurological condition. All experimental procedures were approved by the Research Ethics Committee at Western University. The participants signed a written informed consent before participating in the study and were compensated for their participation.

### Stimuli

Six different consonant-vowel (CV) syllables were visually presented to the participants at the center of the screen using PsychoPy (https://www.psychopy.org/). The syllables were composed of a plosive consonant (/p/, /b/, /t/, /d/, /k/, /g/) followed by the vowel/a/ (Fig. 1A). Plosives are produced by blocking the airflow in the vocal tract and then releasing it, creating a burst of air. The plosive syllables in the current experiment varied along two axes: place of articulation (PoA) and voice onset time (VOT). PoA refers to the location in the vocal tract where the airflow is obstructed. In plosive sounds, the blockage can be formed with the lips (/p/, /b/), with the tongue tip against the alveolar ridge (/t/, /d/), or with the back of the tongue against the velum (/k/, /g/), corresponding to the *bilabial*, *alveolar*, and *velar* sounds, respectively. VOT refers to the interval between the release of the plosive closure and the onset of vocal fold vibration. In *voiceless* plosives (/p/, /t/, /k/), the vocal folds do not vibrate during the release, whereas in *voiced* plosives (/b/, /d/, /g/), the vocal folds do vibrate during the release.

**Figure 1.**
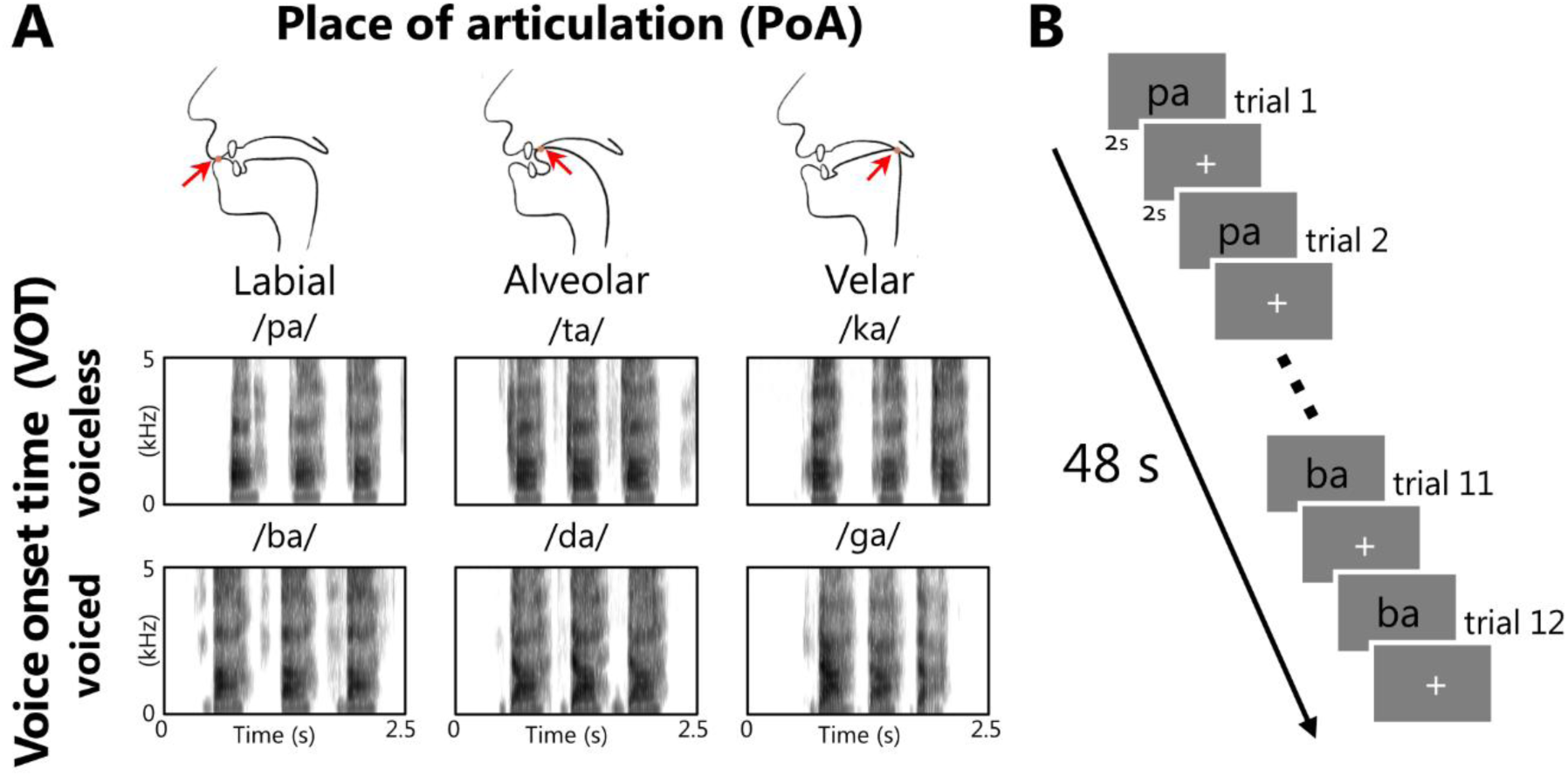
Syllable repetition task. ***A***, Experimental stimuli. Top: Schematic illustration of the vocal tract during articulation of bilabial (/p/,/b/), alveolar (/t/,/d/), and velar (/k/,/g/) plosive consonants. Red arrow denoting the place of articulation. Bottom: Spectrograms of spoken CV syllables of one representative subject (female, 18yo) recorded during the behavioral training session. Spectrograms are grouped by voice onset time, with voiceless plosives (/pa/,/ta/,/ka/) shown in the top row and voiced plosives (/ba/,/da/,/ga/) in the bottom row. ***B***, Block example. Each trial consisted of a CV syllable presented for 2 s, followed by 2 s fixation cross. Participants (N=12) were instructed to repeat the syllable three times during its presentation. Within each block, all six syllables were presented in a random order, with each syllable repeated twice on consecutive trials. Each block lasted ∼48 s. Abbreviations: CV – consonant vowel; PoA – place of articulation; VOT – voice onset time.

### Experimental design

Participants underwent MRI scanning in a single session. We acquired an anatomical image and 10 functional runs. Right before the MRI scan, participants were familiarized with the behavioral task in a short behavioral session (2 runs, ∼20 min). On each trial, a syllable was presented on the screen for 2 seconds, followed by a fixation cross for 2 seconds (Fig. 1B). Participants were instructed to repeat the syllable out load three times at a comfortable pace during the 2s presentation period, and to remain silent during fixation to prevent overlap across task phases. Each run lasted ∼4 minutes and included four blocks of 48 s separated by 14 s rest. Within each block all six syllables were presented, with each syllable appearing twice on consecutive trials. The order of items pairs within a block was randomized, resulting in a total of eight presentations of each item per run. Item and trial repetitions were included to improve the contrast-to-noise ratio (CNR). A period of 10 s rest was added at the end of each functional run to allow for signal relaxation and provide a better estimate of baseline activation. The entire MRI session, including the anatomical scans and setup, lasted ∼60 minutes.

### Imaging data acquisition

Functional MRI data were acquired on a 7T Siemens Magnetom scanner with a 32-channel head coil at Western University. Anatomical T1 weighted scan of each participant was acquired at the beginning of the MRI session, using a magnetization-prepared rapid gradient echo sequence (MP2RAGE, voxel size=0.7mm isotropic; TR=6000 ms; TE=2.27 ms; field of view=246×246; 224 slices). Task-based functional data were acquired using a multi-band gradient-echo EPI sequence with anterior to posterior phase-encoding direction (voxel size= 2.3 mm isotropic; TR=1100 ms; TE=20 ms; flip angle=30; multiband acceleration

Factor=2; GRAPPA acceleration=3; field of view=208×208 mm; 56 slices). Each run consisted of 224 volumes. To correct spatial distortions caused by inhomogeneities in the magnetic field, we also acquired a gradient-echo field map (voxel size=1.3 × 1.3 × 2.5 mm; field of view=210×210).

### Preprocessing

Functional data were preprocessed in native space for each participant separately using SPM12 (fil.ion.ucl.ac.uk/spm) and custom MATLAB code. Our minimal preprocessing pipeline included the following steps: First, functional images were corrected for geometric distortions caused by magnetic field inhomogeneity using the gradient echo field map (Hutton et al., 2002). Then, functional images were realigned to the first volume of the first run to correct for head motion (six parameters: translation x, y, and z, and rotation pitch, roll and yaw). Lastly, the biased-corrected functional data were co-registered to the anatomical T1 image, for which the (0,0,0) coordinate was moved to the anterior commissure (AC). No smoothing or normalization to a group template was performed at this stage.

### First level general linear model

The preprocessed functional images were analyzed with a general linear model (GLM), using a separate regressor for each syllable (/p/, /b/, /t/, /d/, /k/, /g/), for each run. The activation of each trial (consisting of three repetitions of the same syllable) was modeled using a boxcar function of length 2 sec convolved with a two-gamma canonical hemodynamic response function with a peak at 5 sec and a post-stimulus undershoot minimum at 11 seconds. This analysis resulted in activation images (beta maps) for each condition per run, for each participant. Rest was not modeled explicitly but served as an implicit baseline.

### Neocortical surface reconstruction

Reconstruction of cortical surface from the anatomical image was carried out using Freesurfer (Fischl et al., 1999). In this procedure, white-gray matter and pial surfaces were reconstructed for each participant. The surfaces were then inflated into a sphere, and aligned to the left-right symmetric template atlas (fs_LR.32k; Van Essen et al., 2012) based on sulcal depth and curvature information. The functional data were projected from native space to the subject’s individual surface, by averaging beta values of voxels intersecting the line connecting corresponding vertices of the individual white matter and pial surfaces.

### Cerebellar normalization

Cerebellar isolation and segmentation into white and gray matter were performed using the Spatially Unbiased Infratentorial Template (SUIT) toolbox implemented in SPM12 (Diedrichsen, 2006). For each subject, the automatic segmentation was carefully inspected and, when necessary, manually corrected by one of the authors (S.J.) to exclude voxels originating from non-cerebellar tissue (e.g., visual cortex). Cerebellar gray and white matter maps were then normalized into SUIT space using a non-linear deformation algorithm (Ashburner, 2007). The activation estimates (i.e., beta weights) and residual mean-square from the first level GLM were also resliced into SUIT. The functional data were further resliced into MNIsymC template atlas (a symmetric version of the cerebellar only template, aligned to the MNINlin2009cSym template) to enable the definition of symmetric cerebellar ROIs in the left and right hemisphere. For visualization purposes, functional maps were projected onto a flat representation of the cerebellum using the SUIT toolbox (Diedrichsen & Zotow, 2015).

### Defining functional regions of interest

To identify brain regions activated during overt syllable repetition we generated group-level activation maps contrasting the average activation across all syllables against rest. Individual subject data were projected into group space, spatially smoothed on the cortical surface or in the volume (kernel width: 6mm), and averaged across participants. To obtain symmetric ROIs across hemispheres for lateralization analyses, we further averaged the data across hemispheres for ROI definition. Functional ROIs were defined by thresholding the resulting group-average map to retain the top 10% of vertices in the neocortex, and top 10% of voxels in the cerebellum. For cerebellar ROIs, we extracted SPM t-values, reflecting average beta maps divided by the standard deviation of the residual time series at each voxel. For the neo-cortex, anatomical locations of the regions were identified using the Glasser et al. (2016) atlas.

### Quantifying pattern reliability

Pattern reliability was defined as the proportion of total variance in fMRI activity patterns that could be explained by reliable differences between syllables. For each voxel or vertex and each run, we first subtracted the mean response across all syllables. This step removes shared global activations that are not specific to individual syllables. The resulting fMRI activity was decomposed into three variance components: (1) *group*: reflecting patterns that are shared across subjects; (2) *subject*: capturing reliable idiosyncratic differences; and (3) *noise*: representing run-by-run variability of the estimates within each person (https://functional-fusion.readthedocs.io/). These components were estimated from the covariance matrix of activity estimates across voxels, where *x_s_*_,*i*_ denotes the pattern for subject *s*, in run *i*. The average covariance across different people, *cov(x_s,i_, x_t,j_)*, is equal to the group variance; the average covariance within a person across runs, *cov(x_s,i_, x_s,j_)*, is equal to the sum of group and subject variance; and the average total average variance, *var (x_s,i_)*, is equal to all three components. Pattern reliability is then the sum of the group and subject variance components divided by the total variance. Statistical significance of reliability estimates was assessed using one-sample t-tests against zero.

### Group-level univariate analysis

To assess whether mean activation differed systematically as a function of articulatory features, we performed a repeated-measures ANOVA on percent signal change values extracted from each ROI. The model included within-subject factors of Place of Articulation (PoA; bilabial, alveolar, velar), and Voice Onset Time (VOT; voiced, voiceless), with subject treated as a random effect. Significant main effects were followed up with post-hoc comparisons, corrected for multiple comparisons (FDR correction).

To assess the topological organization of different PoA, we computed subject-level contrasts for each articulatory category (bilabial, alveolar, velar) against the remaining syllables. Group-level effects were then estimated by performing a one-sample t-test across participants at each vertex/voxel, yielding group t-statistic maps. These maps were used to evaluate the spatial distribution of place-of-articulation selectivity.

### Multivariate pattern analysis of syllable-specific representations

To quantify how much activation patterns for each syllable differed from each other, we calculated cross-validated distance between pairs of activity patterns (Nili et al., 2014), resulting in a syllable × syllable representational dissimilarity matrix (RDM). Prior to calculating the distances, beta weights for each voxel were divided by the estimated noise standard deviation from the GLM for this voxel. This univariate prewhitening step has been shown to increase the reliability of the RDM (Walther et al., 2016). To obtain a cross-validated estimate of the distances, we multiplied the difference between the activity pattern of two syllables in one imaging run with the difference computed in any other imaging run. This procedure ensures that if two patterns differ only due to noise, then the expected distance estimate is zero (Diedrichsen et al., 2021). For visualization purposes only, the RDM values were square-root transformed and then normalized by their Euclidian norm to remove overall scale differences across ROIs. Note that distances between activity patterns within each ROI are statistically independent of the data used for ROI definition, as ROIs were defined based on average activity across all syllables relative to rest.

### Lateralization index

To assess hemispheric asymmetry during syllable production, we computed a lateralization index for each speech-related ROI. Lateralization index was defined as the normalized difference between right- and left-hemisphere activity (or representational distances) within each ROI. To account for potential negative values, the difference was normalized by the sum of the absolute values of both hemispheres:

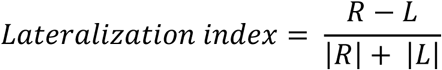

Positive values indicate right-hemisphere dominance, while negative values indicate left-hemisphere dominance. Statistical significance of the lateralization was assessed using one-sample t-tests, testing whether the mean lateralization index differed from zero.

### Comparing representational dissimilarities

To investigate whether the representational structure of syllables differed between hemispheres we calculated the cosine similarity between each subject’s RDM and the leave-one-out group-average RDMs of the left and right hemispheres. This resulted in four similarity measures per subject (per ROI), reflecting how well each hemisphere’s representational structure matched the ipsilateral and contralateral group patterns. Differences between ipsilateral and contralateral similarities were assessed using a paired t-test.

The same method was also used to test whether the superior and inferior cerebellar ROIs differed in their representational structure. For each participant, we computed the cosine similarity between their RDMs and the corresponding leave-one-out group-average RDMs, and compared these similarities using a two-sided paired t-tests.

### Testing of PoA and VOT encoding

To assess whether each phonetic feature (PoA and VOT) was encoded in regional activity patterns, we contrasted three specific types of distances within the RDM. We averaged, for each subject and ROI, RDM entries for pairs with the same PoA (but different VOT, e.g., /pa/ vs. /ba/), the same VOT (but different POA, /pa/ vs. /ta/), or that had none of the features in common (e.g. /pa/ vs. /da/). Encoding of PoA was then measured by the difference between pairs with the same PoA and pairs that had nothing in common. Encoding for VOT was established similarly. This RSA contrast follows the logic of cross-decoding, and tests whether each feature is encoded invariantly across variations in the other feature (Allefeld & Haynes, 2014). Group-level significance was assessed using paired t-test between the two types of differences across participants.

### Model comparison

To compare the relative contribution of each phonetic feature to the overall representation structured we modeled the RDM as a linear combination of three different components: One component that reflects the encoding of the VOT, one component that reflects the encoding of the PoA, and an interaction component (INT). Whereas the VOT and PoA components hypothesize that each feature is encoded independently from other features (Fig. 4C), the interaction component assumes that each feature combination (i.e. each syllable) is encoded by a unique activity pattern. Consequently, the predicted RDM under this model assigns equal dissimilarity to all pairs of distinct syllables. To estimate the contribution of each of these components we performed a non-negative linear regression for each subject, predicting the 15 inter-syllable distances from linear combination of the three predicted RDMs:

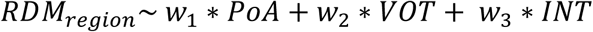

For this analysis, the empirical RDM and the three components (PoA, VOT, and INT) were first vectorized, retaining only the 15 upper triangular elements. These vectors were then standardized to unit norm. The relative contributions of PoA and VOT were then compared using a paired t-test on the regression weights *w*_1_ and *w*_2_ across subjects.

### Multidimensional scaling

To visualize the geometric relationships of syllable representations across speech-related regions, we applied classical multidimensional scaling (MDS) to the vectorized RDM vectors. MDS projects the N-dimensional dissimilarity matrix into a lower dimensional space, while preserving pairwise distances. To focus on the pattern of geometric organization rather than overall magnitude of dissimilarities, each RDM was vectorized and then standardized to unit norm.

### Cerebellum-cortex representational similarity

To quantify the similarity between cerebellar and cortical representations, we computed the cosine similarity between each subject’s cerebellar RDM and each cortical RDM. To assess whether these similarities exceeded what would be expected under a uniform representational structure, we compared each cerebellar-cortical similarity to the INT model, in which all pairwise syllable distances were equal. Differences between cerebellar-cortical and null similarities were evaluated using a one-sided paired t-test.

### Leave-one-out non-negative regression

To identify which cortical areas contribute unique, non-redundant information in explaining cerebellar RDM vectors, we performed a stepwise non-negative linear regression, predicting cerebellar RDMs from the cortical RDMs, by adding one cortical region at each step. The model was trained using data from all but one subject and tested on the left-out subject. Model performance was evaluated using cosine similarity between the predicted and the observed cerebellar RDMs. To assess whether adding a cortical region significantly improved prediction accuracy, we conducted one-sided paired t-test comparing the performance between consecutive steps.

## 3. Results

### 3.1. Identifying regions responsive to syllable production

We identified six neocortical regions and two cerebellar regions activated during overt syllable repetition (Fig. 2A-C). These areas included a large region in the ventral primary motor (M1) and sensorimotor (S1) cortex (vSM). A second, smaller region was identified more dorsally (dSM), situated between the hand and foot representation. We also observed consistent activity in primary auditory cortex and the auditory belt (Aud), the supplementary and pre-supplementary motor areas (SMA), as well as the sylvian parietal-temporal area (Spt). Together, these regions correspond to the “minimal speech production network” described by Bohland & Guenther (2006).

**Figure 2.**
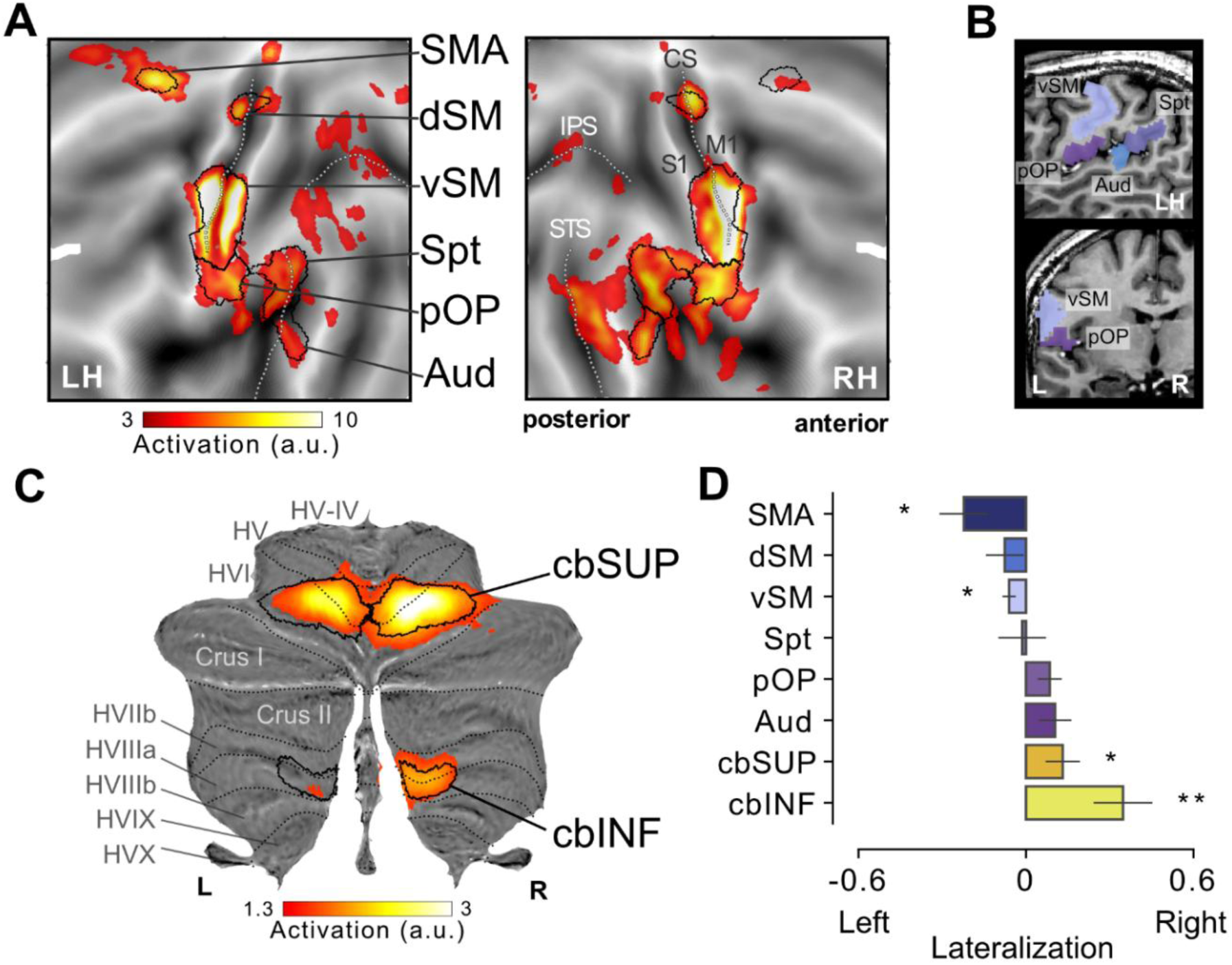
Average activity during overt syllable production. Group-average activation maps across all syllable types, projected onto flattened representation of the neocortex (***A***) and the cerebellum (***C***). Major neocortical sulci and cerebellar lobular boundaries are indicated by dotted black lines. Boundaries of symmetrically-defined functional ROIs are outlined in solid black line. ***B,*** Sagittal (top) and coronal (bottom) sections showing the location of the left pOP in the native space anatomy of a representative subject (male, 22yo). Additional ROIs (vSM, Spt, and Aud) are included for anatomical reference. ***D***, Lateralization index of average activity. Error bar corresponds to standard error of the mean. *p<.05, **p<.01. Abbreviations: SMA- supplementary motor area; dSM – dorsal sensorimotor; vSM – ventral sensorimotor; pOP – parietal operculum; Spt – Sylvian parietal-temporal; Aud – Auditory cortex; CS – central sulcus; S1 – primary sensory cortex; M1 – primary motor cortex; IPS – intraparietal sulcus; STS- superior temporal sulcus; cbSUP – superior cerebellum; cbINF – inferior cerebellum; LH- left hemisphere; L – left; R- right.

Additionally, we identified significant speech-related activity in the parietal operculum. This region was located immediately ventral to vSM (Fig. 2B), yet could be clearly distinguished from it on the cortical flatmap (Fig. 2A). Based on the cytoarchitectonic atlas, the activation was centered primarily in OP4, a subregion of the parietal operculum considered homologous to the parietal ventral area (PV) in non-human primates (Eickhoff et al., 2006).

Within the cerebellum, we identified two distinct regions associated with syllable production: a superior region in lobules V/VI (cbSUP) and an inferior region in lobule VIII (cbINF) (Fig. 2C). This dual representation aligns with previous fMRI findings demonstrating two somatomotor maps for tongue movement in the cerebellum (Nettekoven et al., 2024; Saadon-Grosman et al., 2022).

Speech and language production are typically considered left-lateralized in the cerebral cortex (Hickok & Poeppel, 2007). However, the execution of speech movements engages bilateral sensorimotor cortex (Bohland & Guenther, 2006), suggesting that low-level speech motor control may not follow the same hemispheric specialization observed for higher-level language processes. To test which of the regions within this network show hemispheric asymmetry, we calculate a lateralization index for each region separately (Fig. 2D). We found left hemispheric dominance in the SMA (t_(11)_ =-2.63, p=.023) and vSM (t_(11)_ =-2.97, p=.012). In the cerebellum, both superior and inferior regions showed significantly stronger activity in the right hemisphere (cbSUP: p=.045, cbINF: p=.005).

### 3.2. Pattern analysis shows encoding of different syllables

We then asked whether the identified speech-related regions exhibit distinct activity patterns for different syllables. Surface representations of syllable-related activity patterns showed no clear spatial segregation between syllables (Fig. 3A). Instead, the maps revealed individual differences in both the extent and internal organization of activity patches, consistent with previous fMRI findings (Carey et al., 2017).

**Figure 3.**
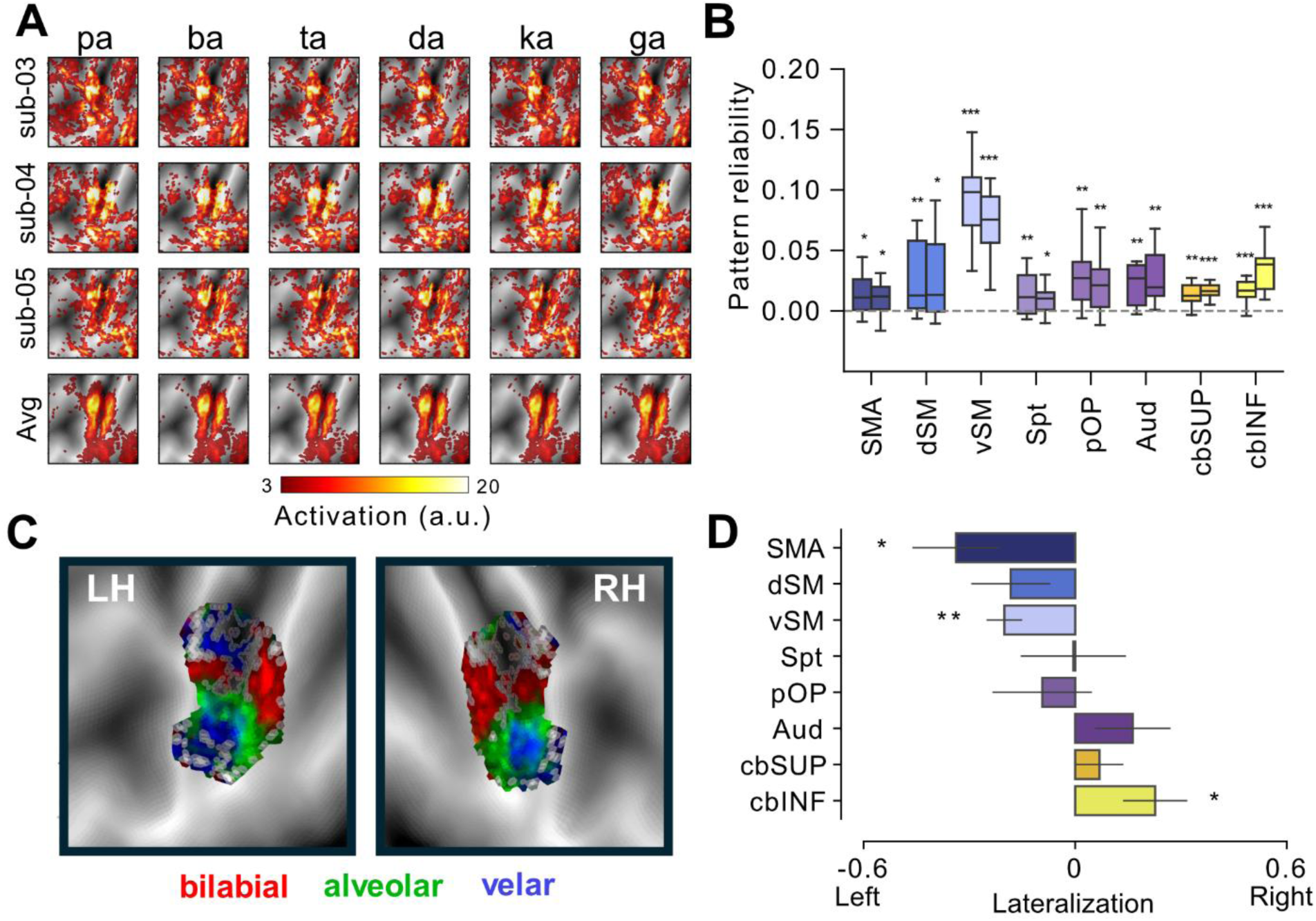
Evoked activity patterns during the production of different syllables. ***A***, Activation maps in the left vSM are shown on a flattened neocortical surface. Each row displays activity patterns from an individual participant, with the bottom row showing the group-average across all subjects (N=12). ***B***, Reliability of activity patterns for single syllables in speech ROIs. Pattern reliability reflects the variance explained by shared variance across subjects and task conditions within each subject, across runs. Within each ROI, reliability for the left hemisphere (left boxplot) and right hemisphere (right boxplot) are presented. ***C***, Group t-maps in left and right vSM showing articulator-selective contrasts for bilabial (red), alveolar (green), and velar (blue). ***D***, Lateralization index calculated as the normalized difference between right and left average Mahalanobis distances, in each ROI. Error bars indicate standard error of the mean. *p<.05, **p<.01, ***p<.001. Abbreviations: SMA- supplementary motor area; dSM – dorsal sensorimotor; vSM – ventral sensorimotor; OP – operculum; Spt – Sylvian parietal-temporal; Aud – Auditory cortex; cbSUP – superior cerebellum; cbINF – inferior cerebellum; LH – left hemisphere; RH – right hemisphere.

To assess whether different syllables were reliably represented despite the absence of clear spatial segregation, we quantified syllable-specific reliability across runs and subjects using variance decomposition (Fig. 3B). Importantly, reliability was estimated after subtracting the mean activity from each voxel, ensuring the measure reflected the consistency of differences between syllables rather than overall activation levels. All ROIs showed significant positive pattern reliability (see *Methods*, all p<0.05, FDR-corrected), indicating that syllable identity could be reliably decoded from these regions in individual subjects.

We then decomposed pattern reliability into two components: group variance, reflecting structure shared across individuals, and subject-specific variance, capturing reliable but idiosyncratic patterns. In vSM, the group component was highly significant (left: t_(11)_=6.79, p_(FDR)_=.0001; right: t_(11)_=5.93, p_(FDR)_=.0001), accounting for 35.93% (±5.19%) of the individual pattern reliability. Thus, despite the apparent lack of common organization on visual inspection (Fig. 3A), vSM exhibits a systematic topology shared across subjects. To illustrate this, we plotted group contrasts for each place of articulation (bilabial, alveolar, velar) against the remaining syllables. The resulting map (Fig. 3C) shows more dorsal activation for bilabials, and more ventral activity for alveolar, with velar syllables engaging voxels in two separate areas, consistent with previous findings (Bouchard et al., 2013; Carey et al., 2017; Correia et al., 2020; Eichert et al., 2020).

Lastly, we tested whether syllable-specific information differed across hemispheres (Fig. 3D). Significant left lateralization was observed in both SMA and vSM, with stronger information in the left hemisphere compared to the right (SMA: t_(11)_=-2.97, p=.012; vSM: t_(11)_ =-4.15, p=.001). The dSM showed a non-significant trend toward left lateralization (t_(11)_ =-1.66, p=.12). In contrast, the inferior cerebellum exhibited a significant right lateralization, with higher information encoded in the right cerebellar hemisphere (t_(11)_=2.55, p=.026). The operculum, auditory cortex, Spt, and superior cerebellum showed no hemispheric differences in the strength of syllable-specific information (all ps >.1).

### 3.3. Representational geometry in cortical speech regions

Having established reliable syllable-specific information, we next examined its organization across regions using representational similarity analysis (Kriegeskorte & Diedrichsen, 2019). For each ROI, we computed the cross-validated Mahalanobis distance between each pair of syllables. A distance estimate greater than zero indicates a reliable difference between the regional activity patterns for these two syllables. All pairwise distance estimates were then combined into a representational dissimilarity matrix (RDM) per region (Fig. 4A).

**Figure 4.**
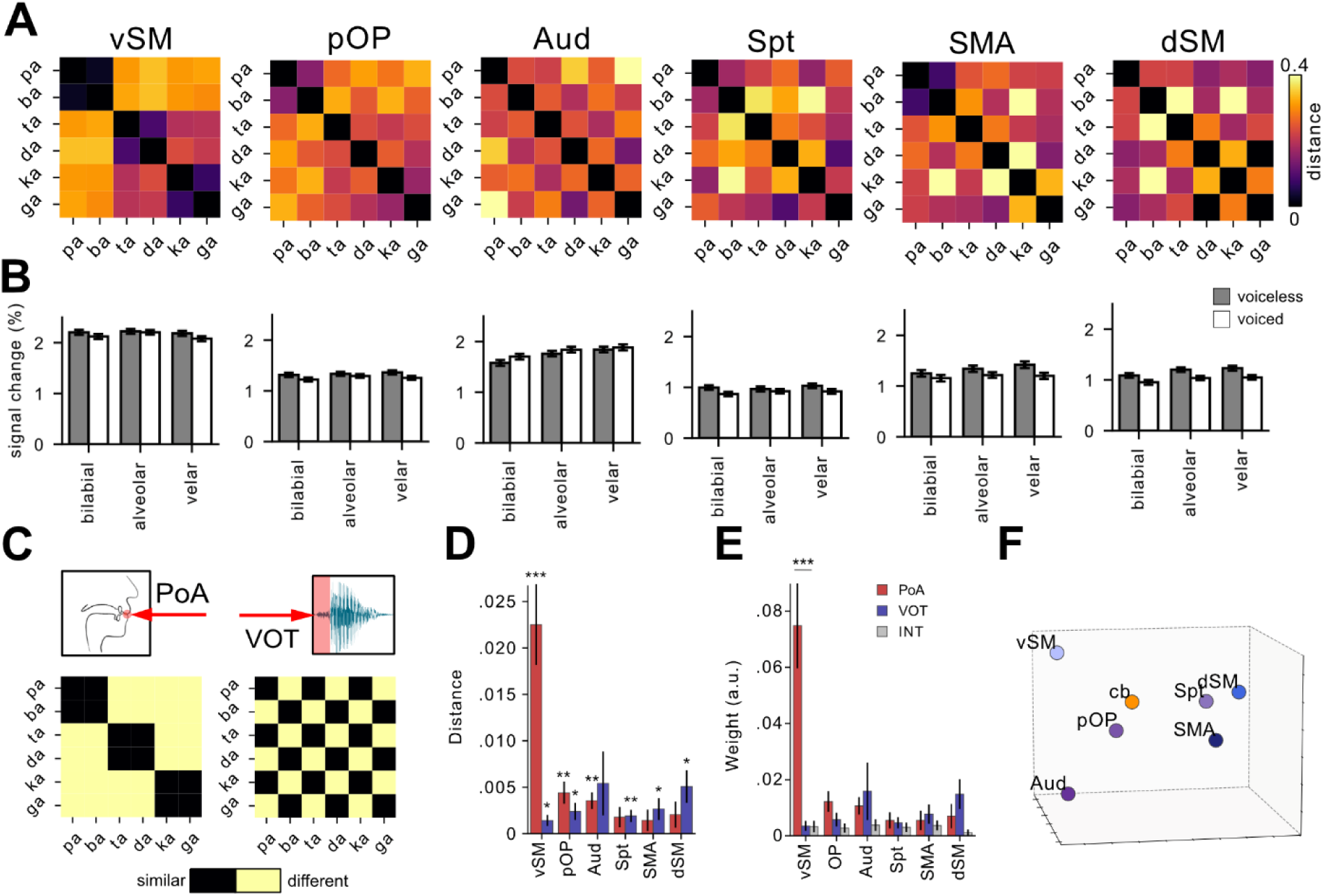
Representational geometry of syllables within the neocortex. ***A,*** RDMs between activity patterns evoked by different syllables, averaged across hemispheres and subjects within each region. ***B***, Percent signal change (mean ± SEM) for each region depicted in A, grouped by POA, with voiced (white) and voiceless (gray) consonants shown separately. ***C***, Distance structure for the PoA (left) and VOT (right) models. Black indicates no difference between syllables; yellow indicates large differences. **D,** Feature isolated neural distances (mean ± SEM) for PoA (red) and VOT (blue) across cortical regions. ***E***, Comparison of syllable model fits across speech related ROIs. Bars show mean weight (*w* ± SEM) across participants for PoA (red), VOT (blue), and INT (gray). ***F,*** Multidimensional scaling of differences between ROIs in three-dimensional space. *p<.05, **p<.01, ***p<.001. Abbreviations: RDM – representational dissimilarity matrix; vSM – ventral sensorimotor; OP –operculum; Aud – Auditory cortex; Spt – Sylvian parietal-temporal; SMA- supplementary motor area; dSM – dorsal sensorimotor; cb-cerebellum; PoA – place of articulation; VOT – voice onset time; INT – interaction.

We first tested whether there were any significant differences in representational geometry between the left and right hemispheric compartments of each functional region. In our task, we found no significant hemispheric differences in the RDMs (see *Methods*, all ps > 0.1). We therefore averaged the data from left and right hemispheres to produce a single representational estimate per ROI. In addition to the pattern differences between syllables, we also quantified mean activation responses for each syllable within each ROI (Fig. 4B).

We then examined how syllables are represented within each region, testing whether the place of articulation (PoA) or voice onset time (VOT) were represented independently from the other feature (Fig. 4D; see *Methods*). This analysis revealed a strong encoding of PoA in the vSM (t_(11)_=5.26, p=6.66×10^-4^), the pOP (t_(11)_=3.97, p=.002), and the auditory cortex (t_(11)_=4.34, p=.001). Univariate analysis revealed significant main effects of PoA in vSM and auditory cortex, consistent with the RSA results. Interestingly, SMA also showed a significant univariate PoA effect, though this was not reflected in the representational geometry (Table 1). Post-hoc comparisons revealed distinct response profiles across these regions: in the vSM alveolar syllables elicited the highest activation, whereas in the SMA and auditory cortex velar syllables evoked the strongest responses (Fig. 4B).

**Table 1.** Repeated-measures ANOVA results for place of articulation (PoA) and voicing (VOT) effects on average activity in each region.

| Region | Factor | F(df1,df2) <sup>a</sup> | p-value | Post-hoc comparison | Post-hoc p-value<sup>b</sup> |
| --- | --- | --- | --- | --- | --- |
| vSM | PoA | F(2,22) = 6.50 | **.0060** | Alveolar > Velar | .020 |
|  | VOT | F(1,11) = 4.73 | .052 |  |  |
|  | PoA × VOT | F(2,22) = 1.00 | .383 |  |  |
| pOP | PoA | F(2,22) = 1.69 | .206 |  |  |
|  | VOT | F(1,11) = 6.87 | **.023** | Voiceless > Voiced | .023 |
|  | PoA × VOT | F(2,22) = 0.69 | .510 |  |  |
| Aud | PoA | F(2,22) = 24.86 | **< .001** | Bilabial < Alveolar<br>Bilabial < Velar | .001<br><.001 |
|  | VOT | F(1,11) = 1.92 | .193 |  |  |
|  | PoA × VOT | F(2,22) = 0.44 | .650 |  |  |
| Spt | PoA | F(2,22) = 1.11 | .348 |  |  |
|  | VOT | F(1,11) = 9.75 | **.0097** | Voiceless > Voiced | .0097 |
|  | PoA × VOT | F(2,22) = 0.75 | .485 |  |  |
| SMA | PoA | F(2,22) = 4.91 | **.017** | Bilabial < Velar | .011 |
|  | VOT | F(1,11) = 12.32 | **.0049** | Voiceless > Voiced | .0049 |
|  | PoA × VOT | F(2,22) = 1.55 | .235 |  |  |
| dSM | PoA | F(2,22) = 6.36 | **.0066** | Bilabial > Velar | .025 |
|  | VOT | F(1,11) = 17.29 | **.0016** | Voiceless > Voiced | .0016 |
|  | PoA × VOT | F(2,22) = 0.19 | .832 |  |  |
| cb | PoA | F(2,22) = 10.35 | **< .001** | Bilabial < Alveolar<br>Bilabial < Velar | .000367<br>.020 |
|  | VOT | F(1,11) = 3.73 | .079 |  |  |
|  | PoA × VOT | F(2,22) = 0.89 | .423 |  |  |

<sup>a</sup>F values are reported with numerator and denominator degrees of freedom.
<sup>b</sup>Post hoc p values are FDR-corrected.

Encoding of VOT was observed in dSM (t_(11)_=3.02, p=.011), SMA (t_(11)_=2.43, p=.033), Spt (t_(11)_=3.34, p=.006), pOP (t_(11)_=2.89, p=.014), and vSM (t_(11)_=2.82, p=.016), indicating sensitivity to voicing structure in these regions. Consistent with these representational effects, univariate analysis showed significant main effects of VOT in pOP, Spt, SMA and dSM (Table 1), with voiceless syllables eliciting stronger responses across all regions (Fig. 4B).

Next, we compared the contribution of each feature in explaining the regional RDMs (Fig. 4E). We used non-negative linear regression to simultaneously fit each cortical RDM with the two model RDMs, along with an interaction term capturing unique syllable-specific patterns. We then compared the resulting feature weights within each region. vSM showed a significantly stronger weight for PoA compared to VOT (t_(11)_=4.85, p=.0005). The parietal operculum also exhibited a slight preference for PoA over VOT, albeit not significant (t_(11)_=2.15, p=.054). Other cortical regions showed no significant difference between PoA and VOT weights (all ps>.1), with the auditory cortex, SMA and dSM showing a trend towards stronger VOT encoding.

Multidimensional scaling (MDS; Fig. 4F) visualizes the organizational structure across cortical regions. Regions sensitive primarily to PoA (vSM and auditory cortex) were spatially separated from those showing VOT sensitivity (dSM and SMA). The spatial position of pOP between these groups is consistent with its sensitivity to both phonetic features.

### 3.4. Representational geometry in the cerebellum

We next investigated whether the superior and inferior cerebellar speech regions differed in their representational structures for speech features. Contrary to this hypothesis, comparison of the representational geometry between the two regions revealed no significant difference. The RDM of the superior region of an individual participant was equally similar to the group average of the superior and inferior cerebellar regions (p>0.3; Fig. 5A). Despite their anatomical segregation, the two cerebellar speech zones therefore appear to organize syllables according to a shared representational structure. Accordingly, we averaged the RDMs across these two cerebellar regions to obtain a more reliable, consolidated measure of syllable organization within the cerebellum (Fig. 5B). Mean activations for each condition were also averaged across these regions to complement the representational analysis with univariate measures (Fig. 5C).

**Figure 5.**
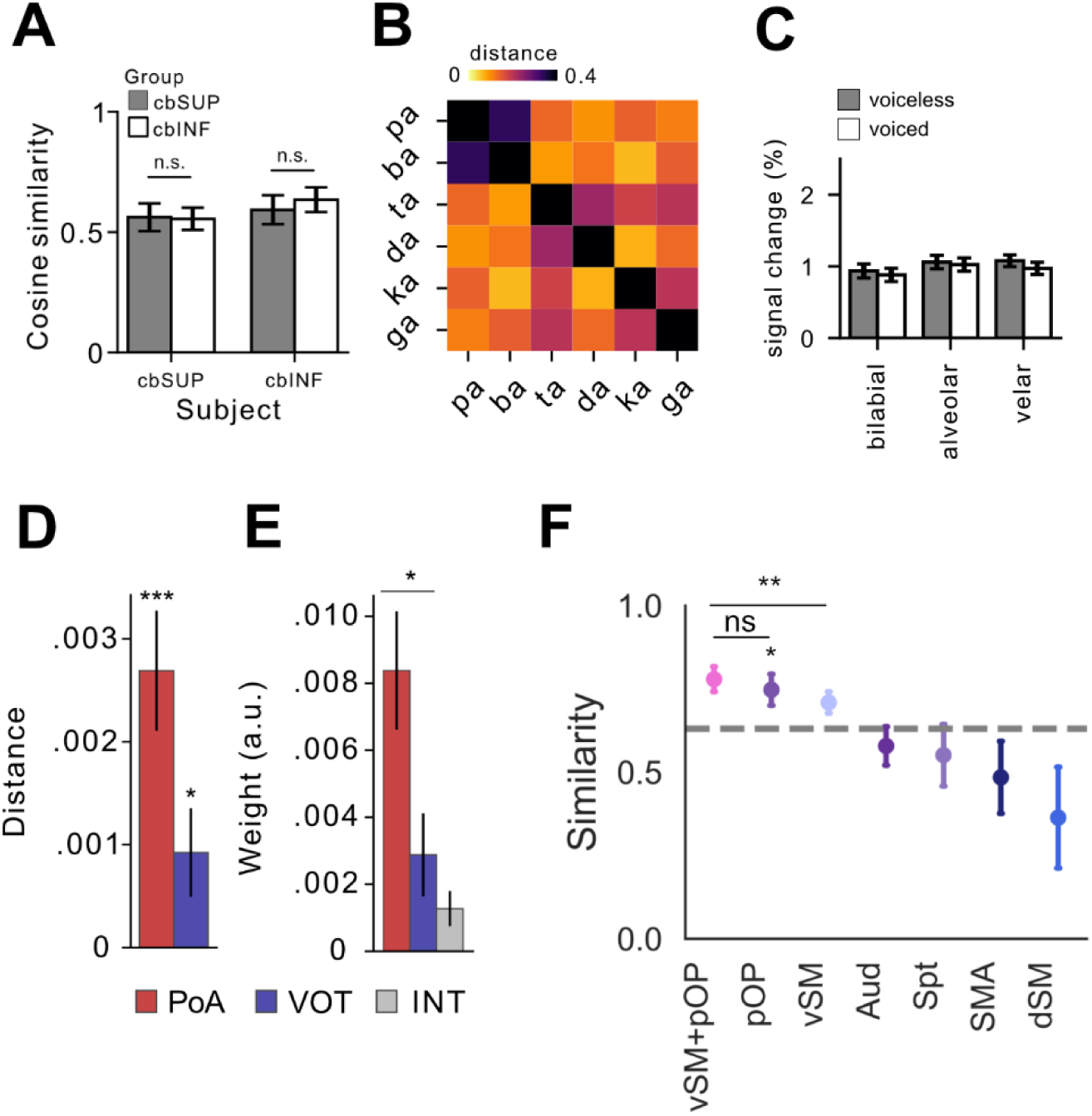
Representational geometry of syllables within the cerebellum. ***A,*** Cosine similarities between cerebellar ROIs. Plotted are the cosine similarities of each participant’s cbSUP and cbINF RDMs compared to the leave-one-out group RDMs of cbSUP (gray) and cbINF (white). ***B***, RDMs between activity patterns evoked by different syllables, averaged across cbSUP and cbINF. ***C***, Percent signal change (mean ± SEM), grouped by POA, with voiced (white) and voiceless (gray) consonants shown separately. ***D*,** Feature isolated neural distances (mean ± SEM) for PoA (red) and VOT (blue). ***E***, Size of model weights (*w* ± SEM) across participants for PoA (red), VOT (blue), and INT (gray) in the combined model. ***F***, Cosine similarity between the cerebellum and neocortex. Similarities were calculated between observed and predicted cerebellar RDMs using cross validated stepwise non-negative regression. The dashed gray line represents the similarity of cerebellar RDMs to the INT model, averaged across participants. *p<.05, **p<.01, ***p<.001. Abbreviations: RDM – representational dissimilarity matrix; cbINF- inferior cerebellum; cbSUP- superior cerebellum; PoA – place of articulation; VOT – voice onset time; INT – interaction; vSM – ventral sensorimotor; OP –operculum; Aud – Auditory cortex; Spt – Sylvian parietal-temporal; SMA- supplementary motor area; dSM – dorsal sensorimotor.

To characterize the geometrical structure of cerebellar syllable representations, we tested their relationships with each of the phonetic features. This analysis revealed both significant encoding of PoA (t_(11)_=4.70, p=.0006), and of VOT (t_(11)_=2.21, p=.048), indicating that the cerebellum encodes both articulatory structure and voice onset (Fig. 5D). Univariate analysis was consistent with the PoA findings, showing a main effect of PoA with bilabial sounds eliciting lower activation than other place categories (Table 1). When comparing the strength of the two features using the weights from non-negative linear regression, we found that PoA was encoded more strongly than VOT (t_(11)_=2.55, p=.026) (Fig. 5E).

The arrangement of representations in speech regions (Fig. 4F) suggests that the representational structure in cerebellar speech areas lies somewhere between vSM and dSM and actually most closely resembles that of the parietal operculum. To quantify this observation, we calculated cosine similarity between the cerebellum and each cortical region. Because of substantial inter-individual variability in cerebellar representational structure, this analysis was performed at the individual-subject level to capture subject-specific relationships between cortical and cerebellar representations. Within each subject, the cerebellar RDM was significantly more similar to the pOP than to INT model (t_(11)_=2.206, p=.025 (Fig. 5F). Cerebellar RDMs also showed some similarity to vSM, though this did not reach statistical significance (p=.059). Similarities to other regions were not significant, indicating a selective engagement of the cerebellum with the pOP and vSM during speech production.

To determine whether cerebellar speech representations reflect coupling with a single cortical region or a mixture of cortical sources, we estimated weights that best predicted cerebellar RDMs based on cortical RDMs, using a leave-one-out cross-validation approach incrementally adding ROIs based on their similarity to the cerebellar RDM. This analysis revealed that a single region, the pOP, was sufficient to account for cerebellar syllable representations. Adding the pOP to a model already including vSM significantly improved prediction of cerebellar RDMs (t_(11)_=2.65, p=.002), whereas adding vSM to a model that already included the pOP did not (p=0.1). Overall, these results suggest that the cerebellar speech regions are most similar to pOP, rather than reflecting a mixture of cortical representations (Fig. 5F).

## 4. Discussion

Using 7T fMRI and multivariate representational analysis, we examined how the brain encodes different features of syllable production, place of articulation and voice onset time, across cortical and cerebellar regions. Within primary sensorimotor cortex, we found evidence of a functional dissociation: ventral regions encoded syllables primarily according to articulatory configuration, while dorsal regions showed a greater sensitivity to the temporal onset of voicing. In addition to other expected areas, we identified a region in the parietal operculum that encoded a combination of place of articulation and phonation. Addressing our main research question, we found that both the superior and inferior cerebellar speech regions exhibited a similar mixed representation of these two speech features. Rather than mirroring the representational structure of either the ventral or dorsal primary sensorimotor speech region, the cerebellar representations aligned most closely with those of the parietal operculum. These findings suggest an opercular-cerebellar circuit involved in speech motor control that operates beyond the level of primary motor execution.

### 4.1. Speech representations in the primary motor cortex

In this study, we found a functional contrast between vSM and dSM in their tuning to speech features. The tuning of the vSM for place of articulation, and its somatotopic organization, are consistent with its established role in the control of articulatory muscles (Bouchard et al., 2013; Carey et al., 2017). While dorsal sensorimotor cortex has traditionally been associated with trunk movements involved in voluntary breathing (Foerster, 1931; Penfield & Boldrey, 1937), our current findings add representation-level evidence that its role in speech motor control extends beyond respiration. Correia et al. (2020) first suggested this idea based on greater activation in dSM during voiced versus whispered speech that persisted even when respiratory demands were matched. Here, we extended this by showing that dSM represent syllables according to their voice onset time, demonstrating structured encoding of voicing distinctions across multiple place-of-articulation contexts. Together, these findings support a functional dissociation within primary sensorimotor cortex: vSM encodes articulatory configuration organized somatotopically along the vocal tract, while dSM encodes characteristics of voicing.

It should be noted that the VOT sensitivity in the dSM did not significantly dominate PoA in a non-negative linear regression model, suggesting that dSM is not exclusively tuned to voicing. One possible explanation is that dSM represents multiple effectors, with laryngeal representations required for phonation is embedded alongside representations of other body parts. Alternatively, VOT sensitivity in dSM may partly reflect differences in respiratory motor demands rather than phonation per se. Voiceless plosives typically require greater subglottal pressure than voiced plosives, which in turn may involve greater recruitment of respiratory and thoracic musculature. If dSM contains representations related to respiratory and thoracic musculature, increased activity for voiceless sounds could reflect these additional motor demands rather than selective encoding of voicing. Because the present study did not control respiratory demands across voiced and voiceless conditions, future studies will be needed to dissociate phonatory from respiratory contributions to dSM activity.

### 4.2. Parietal operculum

Our data revealed a speech-related area in the parietal operculum that was anatomically separate from the adjacent vSM (Fig. 2A-B) and showed a distinct representational organization encoding both place of articulation and voice onset time (Fig. 4). The parietal operculum has been subdivided into four cytoarchitectonically and functionally defined areas (OP1-4; Burton et al., 2008; Eickhoff, Schleicher, et al., 2006), with OP1 and OP4 corresponding to secondary somatosensory cortex (SII) parietal ventral area (PV) in non-human primates, respectively (Eickhoff, Amunts, et al., 2006). In humans, OP1 has been mainly implicated in light touch, pain, and tactile discrimination (Burton et al., 2008). Speech related activity in the present study, however, was mostly observed in OP4, which is functionally interconnected at rest with auditory and motor speech-related areas (Sepulcre, 2015). The structured encoding of place of articulation and voice onset time observe in OP4 further supports a role for this area in speech production.

Previous fMRI studies of speech production have reported speech-related activity in the approximate vicinity of OP4, but its interpretation has been inconsistent. This activity has been variously ignored or attributed to the Rolandic operculum (Bohland & Guenther, 2006; Tourville et al., 2008), or parietal operculum (Bailey et al., 2021), reflecting inconsistencies in anatomical nomenclature across the literature. Precise localization is further complicated by the proximity of OP4 to the posterior insula and superior temporal cortex, regions that can be difficult to distinguish in volume-based analyses. By defining ROIs on the cortical surface and anchoring them to contemporary cytoarchitectonic parcellations, the present study localizes speech-related representations to OP4 with greater anatomical precision.

### 4.3. Cerebellar speech representations

Our main goal was to determine how cerebellar speech representations fit into the neocortical speech network. We found that the cerebellum encodes both place of articulation and voicing, with a stronger representation of place of articulation. This pattern is consistent with the proposed role of the cerebellum in coordinating articulatory movements during speech, as well as with clinical observations of articulatory impairments following cerebellar damage (Ackermann et al., 1992). Representations of both articulatory and phonatory speech features have previously been reported in the cerebellum (Correia et al. 2020). The present findings extend this work by directly comparing the relative contribution of these dimensions, revealing that place of articulation contributes significantly more to the representational structure than voicing.

However, articulatory feature dominance in the cerebellum was substantially weaker than in vSM. Accordingly, cerebellar syllable representations were not best explained by the representational structure observed in vSM. This finding is difficult to reconcile with a simple closed-loop cortico-cerebellar motor model, in which each cerebellar motor territory is reciprocally connected with a single corresponding sensorimotor cortical region (Kelly & Strick, 2003; Saadon-Grosman et al., 2022). While this framework appears to capture the relationship between cortical and cerebellar representations of hand movements (King et al., 2023; Wiestler et al., 2011), our results suggest that it does not generalize to speech production. Instead, cerebellar speech representations were most similar to those observed in the parietal operculum.

We propose several possible explanations for this finding. First, because cerebellar BOLD signals are thought to predominantly reflect inputs to the cerebellar cortex rather than the output of Purkinje cells (Caesar et al., 2003; Thomsen et al., 2009), the similarity between cerebellar and opercular representational geometries may indicate that the cerebellar speech regions receive a substantial portion of its input from the parietal operculum. Alternatively, the cerebellum may integrate inputs from multiple speech-related motor regions, including vSM, dSM and potentially other cortical areas. Either way, the integrated representation of speech features in the cerebellum could support the role of these areas in the coordination of articulation and phonation.

Previous studies have reported functional differences between superior and inferior cerebellar speech regions, with activity varying as a function of task complexity and timing demands (Bohland & Guenther, 2006; Riecker et al., 2000). In contrast, we found no evidence that these regions differed in their representational organization of speech. This finding is consistent with previous reports of dual representations of body parts within sensorimotor regions of the cerebellum, despite a lack of clear functional differentiation between these spatially segregated representations (Nettekoven et al., 2024; Wiestler et al., 2011). While our current data suggest that there is no difference in the level of phonetic feature encoding between these regions, future studies are required to evaluate their functional profiles across a wider spectrum of conditions.

### 4.4. Temporal-parietal speech representations

Speech production relies on interactions between the motor and auditory systems (Guenther, 2006; Hickok et al., 2011). Our results show that speech evokes robust activations in temporal and parietal regions, including the auditory cortex and the Sylvian parietal-temporal area (Spt), that are implicated in translating auditory feedback into motor commands during speech production (Buchsbaum et al., 2001). Previous studies have shown that the auditory cortex encodes acoustic-phonetic features during speech perception (Binder et al., 2000; Mesgarani et al., 2014). Our data extends these findings by demonstrating that the auditory cortex represents phonatory features not only during speech perception, but also during speech production. Spt, on the other hand, did not show a clear representational structure of syllables, despite showing stronger mean activation for voiceless than voiced sounds. What drives this univariate sensitivity without corresponding representational structure remains unclear.

### 4.5. Lateralization of speech representations

In line with the left-dominant organization of speech motor systems (Hickok & Poeppel, 2007), we observed left lateralization of both mean activation and syllable-specific representational information in vSM and SMA. This extends prior findings to the level of fine-grained representational structure, suggesting that the left hemisphere not only activates more strongly during speech but also represents phonetic features more distinctly. It should be noted that our current experimental design does not allow us to dissociate motor preparation from execution. The left lateralization in vSM and SMA may therefore reflect motor preparation, execution, or both.

In the cerebellum, the pattern was reversed: both superior and inferior regions showed right lateralization of mean activation, consistent with prior reports of right cerebellar dominance during speech production (Bohland & Guenther, 2006; Correia et al., 2020). Given the crossed organization of cortico-cerebellar projections, right cerebellar dominance is the expected counterpart of left cortical dominance, and our findings extend this observation to the level of representational information.

### 4.6. Summary and conclusions

In summary, we found a functional dissociation within primary sensorimotor cortex, with ventral regions encoding place of articulation and dorsal regions encoding voice onset time. The parietal operculum, in contrast, encoded both phonetic dimensions, suggesting an integrative representation of speech features. Contrary to predictions derived from simple cortico-cerebellar motor circuit models, cerebellar speech representations did not mirror those of primary sensorimotor cortex. Instead, they were most closely aligned with representations in the parietal operculum, a higher-order somatosensory region situated at the interface between auditory and motor speech systems. These findings point to an opercular-cerebellar circuit as a key component of the speech network and suggest that the cerebellum contributes to higher-order sensorimotor computations that extend beyond the control of articulatory movements alone.

## Data and code availability

All code used in this paper is available at https://github.com/sjossinger/Articulotopy. The raw MRI data used in this paper is available upon request.

## Author contributions

S.J. contributed to conceptualization, data curation, formal analysis, investigation, visualization, writing – original draft, writing – review and editing. B.A. contributed to data curation, writing – review and editing. J.D. contributed to conceptualization, funding acquisition, investigation, supervision, writing – review and editing.

## References

Ackermann, H. (2008). Cerebellar contributions to speech production and speech perception: Psycholinguistic and neurobiological perspectives. Trends in Neurosciences, 31(6), 265–272. 10.1016/j.tins.2008.02.011

Ackermann, H., & Brendel, B. (2016). Neurobiology of Speech Production: A Motor Control Perspective. In Chapter 59—Neurobiology of Language (pp. 741–750). Academic Press. 10.1016/B978-0-12-407794-2.00059-6

Ackermann, H., & Hertrich, I. (1997). Voice onset time in ataxic dysarthria. Brain and Language, 56(3), 321–333. 10.1006/brln.1997.1740

Ackermann, H., Hertrich, I., Daum, I., Scharf, G., & Spieker, S. (1997). Kinematic analysis of articulatory movements in central motor disorders. Movement Disorders, 12(6), 1019–1027. 10.1002/mds.870120628

Ackermann, H., Hertrich, I., & Scharf, G. (1995). Kinematic Analysis of Lower Lip Movements in Ataxic Dysarthria. Journal of Speech, Language, and Hearing Research, 38(6), 1252–1259. 10.1044/jshr.3806.1252

Ackermann, H., Vogel, M., Petersen, D., & Poremba, M. (1992). Speech deficits in ischaemic cerebellar lesions. Journal of Neurology, 239(4), 223–227. 10.1007/BF00839144

Allefeld, C., & Haynes, J.-D. (2014). Searchlight-based multi-voxel pattern analysis of fMRI by cross-validated MANOVA. NeuroImage, 89, 345–357. 10.1016/j.neuroimage.2013.11.043

Ashburner, J. (2007). A fast diffeomorphic image registration algorithm. NeuroImage, 38(1), 95–113. 10.1016/j.neuroimage.2007.07.007

Bailey, L. M., Bodner, G. E., Matheson, H. E., Stewart, B. M., Roddick, K., O’Neil, K., Simmons, M., Lambert, A. M., Krigolson, O. E., Newman, A. J., & Fawcett, J. M. (2021). Neural correlates of the production effect: An fMRI study. Brain and Cognition, 152, 105757. 10.1016/j.bandc.2021.105757

Binder, J. R., Frost, J. A., Hammeke, T. A., Bellgowan, P. S., Springer, J. A., Kaufman, J. N., & Possing, E. T. (2000). Human temporal lobe activation by speech and nonspeech sounds. Cerebral Cortex (New York, N.Y.: 1991), 10(5), 512–528. 10.1093/cercor/10.5.512

Bohland, J. W., & Guenther, F. H. (2006). An fMRI investigation of syllable sequence production. NeuroImage, 32(2), 821–841. 10.1016/j.neuroimage.2006.04.173

Bouchard, K. E., Mesgarani, N., Johnson, K., & Chang, E. F. (2013). Functional organization of human sensorimotor cortex for speech articulation. Nature, 495(7441), 327–332. 10.1038/nature11911

Buchsbaum, B. R., Hickok, G., & Humphries, C. (2001). Role of left posterior superior temporal gyrus in phonological processing for speech perception and production. Cognitive Science, 25(5), 663–678. 10.1207/s15516709cog2505_2

Burton, H., Sinclair, R. J., Wingert, J. R., & Dierker, D. L. (2008). Multiple parietal operculum subdivisions in humans: Tactile activation maps. Somatosensory & Motor Research, 25(3), 149–162. 10.1080/08990220802249275

Caesar, K., Gold, L., & Lauritzen, M. (2003). Context sensitivity of activity-dependent increases in cerebral blood flow. Proceedings of the National Academy of Sciences, 100(7), 4239–4244. 10.1073/pnas.0635075100

Carey, D., Krishnan, S., Callaghan, M. F., Sereno, M. I., & Dick, F. (2017). Functional and Quantitative MRI Mapping of Somatomotor Representations of Human Supralaryngeal Vocal Tract. Cerebral Cortex, cercor;bhw393v2. 10.1093/cercor/bhw393

Chen, S. H. A., & Desmond, J. E. (2005). Cerebrocerebellar networks during articulatory rehearsal and verbal working memory tasks. NeuroImage, 24(2), 332–338. 10.1016/j.neuroimage.2004.08.032

Correia, J. M., Caballero-Gaudes, C., Guediche, S., & Carreiras, M. (2020). Phonatory and articulatory representations of speech production in cortical and subcortical fMRI responses. Scientific Reports, 10(1), 4529. 10.1038/s41598-020-61435-y

Diedrichsen, J. (2006). A spatially unbiased atlas template of the human cerebellum. NeuroImage, 33(1), 127–138. 10.1016/j.neuroimage.2006.05.056

Diedrichsen, J., Berlot, E., Mur, M., Schütt, H. H., Shahbazi, M., & Kriegeskorte, N. (2021). Comparing representational geometries using whitened unbiased-distance-matrix similarity (arXiv:2007.02789). arXiv. 10.48550/arXiv.2007.02789

Diedrichsen, J., & Zotow, E. (2015). Surface-Based Display of Volume-Averaged Cerebellar Imaging Data. PLOS ONE, 10(7), e0133402. 10.1371/journal.pone.0133402

Eichert, N., Papp, D., Mars, R. B., & Watkins, K. E. (2020). Mapping Human Laryngeal Motor Cortex during Vocalization. Cerebral Cortex, 30(12), 6254–6269. 10.1093/cercor/bhaa182

Eickhoff, S. B., Schleicher, A., Zilles, K., & Amunts, K. (2006). The Human Parietal Operculum. I. Cytoarchitectonic Mapping of Subdivisions. Cerebral Cortex, 16(2), 254–267. 10.1093/cercor/bhi105

Fischl, B., Sereno, M. I., Tootell, R. B. H., & Dale, A. M. (1999). High-resolution intersubject averaging and a coordinate system for the cortical surface. Human Brain Mapping, 8(4), 272–284. 10.1002/(SICI)1097-0193(1999)8:4<272::AID-HBM10>3.0.CO;2-4

Foerster. (1931). The cerebral cortex in man. The Lancet, 218(5632), 309–312. 10.1016/S0140-6736(00)47063-7

Glasser, M. F., Coalson, T. S., Robinson, E. C., Hacker, C. D., Harwell, J., Yacoub, E., Ugurbil, K., Andersson, J., Beckmann, C. F., Jenkinson, M., Smith, S. M., & Van Essen, D. C. (2016). A multi-modal parcellation of human cerebral cortex. Nature, 536(7615), 171–178. 10.1038/nature18933

Gordon, E. M., Chauvin, R. J., Van, A. N., Rajesh, A., Nielsen, A., Newbold, D. J., Lynch, C. J., Seider, N. A., Krimmel, S. R., Scheidter, K. M., Monk, J., Miller, R. L., Metoki, A., Montez, D. F., Zheng, A., Elbau, I., Madison, T., Nishino, T., Myers, M. J.,… Dosenbach, N. U. F. (2023). A somato-cognitive action network alternates with effector regions in motor cortex. Nature, 617(7960), 351–359. 10.1038/s41586-023-05964-2

Guenther, F. H. (2006). Cortical interactions underlying the production of speech sounds. Journal of Communication Disorders, ASHA 2005 Research Symposium: Physiological Foundations of Speech Motor Development and Production, 39(5), 350–365. 10.1016/j.jcomdis.2006.06.013

Guenther, F. H., & Hickok, G. (2016). Chapter 58—Neural Models of Motor Speech Control. In G. Hickok & S. L. Small (Eds.), Neurobiology of Language (pp. 725–740). Academic Press. 10.1016/B978-0-12-407794-2.00058-4

Hickok, G., Houde, J., & Rong, F. (2011). Sensorimotor Integration in Speech Processing: Computational Basis and Neural Organization. Neuron, 69(3), 407–422. 10.1016/j.neuron.2011.01.019

Hickok, G., & Poeppel, D. (2007). The cortical organization of speech processing. Nature Reviews Neuroscience, 8(5), 393–402. 10.1038/nrn2113

Hutton, C., Bork, A., Josephs, O., Deichmann, R., Ashburner, J., & Turner, R. (2002). Image Distortion Correction in fMRI: A Quantitative Evaluation. NeuroImage, 16(1), 217–240. 10.1006/nimg.2001.1054

Ito, M. (2008). Control of mental activities by internal models in the cerebellum. Nature Reviews Neuroscience, 9(4), 304–313. 10.1038/nrn2332

Jürgens, U. (2002). Neural pathways underlying vocal control. Neuroscience and Biobehavioral Reviews, 26(2), 235–258. 10.1016/s0149-7634(01)00068-9

Kelly, R. M., & Strick, P. L. (2003). Cerebellar Loops with Motor Cortex and Prefrontal Cortex of a Nonhuman Primate. Journal of Neuroscience, 23(23), 8432–8444. 10.1523/JNEUROSCI.23-23-08432.2003

King, M., Shahshahani, L., Ivry, R. B., & Diedrichsen, J. (2023). A task-general connectivity model reveals variation in convergence of cortical inputs to functional regions of the cerebellum. eLife, 12, e81511. 10.7554/eLife.81511

Kriegeskorte, N., & Diedrichsen, J. (2019). Peeling the Onion of Brain Representations. Annual Review of Neuroscience, 42, 407–432. 10.1146/annurev-neuro-080317-061906

Mesgarani, N., Cheung, C., Johnson, K., & Chang, E. F. (2014). Phonetic Feature Encoding in Human Superior Temporal Gyrus. Science, 343(6174), 1006–1010. 10.1126/science.1245994

Nettekoven, C., Zhi, D., Shahshahani, L., Pinho, A. L., Saadon-Grosman, N., Buckner, R. L., & Diedrichsen, J. (2024). A hierarchical atlas of the human cerebellum for functional precision mapping. Nature Communications, 15(1), 8376. 10.1038/s41467-024-52371-w

Nili, H., Wingfield, C., Walther, A., Su, L., Marslen-Wilson, W., & Kriegeskorte, N. (2014). A Toolbox for Representational Similarity Analysis. PLOS Computational Biology, 10(4), e1003553. 10.1371/journal.pcbi.1003553

Oldfield, R. C. (1971). The assessment and analysis of handedness: The Edinburgh inventory. Neuropsychologia, 9(1), 97–113. 10.1016/0028-3932(71)90067-4

Penfield, W., & Boldrey, E. (1937). Somatic motor and sensory representation in the cerebral cortex of man as studied by electrical stimulation. Brain, 60(4), 389–443. 10.1093/brain/60.4.389

Riecker, A., Ackermann, H., Wildgruber, D., Meyer, J., Dogil, G., Haider, H., & Grodd, W. (2000). Articulatory/Phonetic Sequencing at the Level of the Anterior Perisylvian Cortex: A Functional Magnetic Resonance Imaging (fMRI) Study. Brain and Language, 75(2), 259–276. 10.1006/brln.2000.2356

Saadon-Grosman, N., Angeli, P. A., DiNicola, L. M., & Buckner, R. L. (2022). A third somatomotor representation in the human cerebellum. Journal of Neurophysiology, 128(4), 1051–1073. 10.1152/jn.00165.2022

Sepulcre, J. (2015). An OP4 Functional Stream in the Language-Related Neuroarchitecture. Cerebral Cortex, 25(3), 658–666. 10.1093/cercor/bht256

Thomsen, K., Piilgaard, H., Gjedde, A., Bonvento, G., & Lauritzen, M. (2009). Principal Cell Spiking, Postsynaptic Excitation, and Oxygen Consumption in the Rat Cerebellar Cortex. Journal of Neurophysiology, 102(3), 1503–1512. 10.1152/jn.00289.2009

Tourville, J. A., Reilly, K. J., & Guenther, F. H. (2008). Neural mechanisms underlying auditory feedback control of speech. NeuroImage, 39(3), 1429–1443. 10.1016/j.neuroimage.2007.09.054

Van Essen, D. C., Glasser, M. F., Dierker, D. L., Harwell, J., & Coalson, T. (2012). Parcellations and Hemispheric Asymmetries of Human Cerebral Cortex Analyzed on Surface-Based Atlases. Cerebral Cortex, 22(10), 2241–2262. 10.1093/cercor/bhr291

Walther, A., Nili, H., Ejaz, N., Alink, A., Kriegeskorte, N., & Diedrichsen, J. (2016). Reliability of dissimilarity measures for multi-voxel pattern analysis. NeuroImage, 137, 188–200. 10.1016/j.neuroimage.2015.12.012

Wiestler, T., McGonigle, D. J., & Diedrichsen, J. (2011). Integration of sensory and motor representations of single fingers in the human cerebellum. Journal of Neurophysiology, 105(6), 3042–3053. 10.1152/jn.00106.2011

Wolpert, D. M., Miall, R. C., & Kawato, M. (1998). Internal models in the cerebellum. Trends in Cognitive Sciences, 2(9), 338–347. 10.1016/S1364-6613(98)01221-2

